# Simple Plans or Sophisticated Habits? State, Transition and Learning Interactions in the Two-step Task

**DOI:** 10.1101/021428

**Authors:** Thomas Akam, Rui Costa, Peter Dayan

**Affiliations:** Champalimaud Neuroscience Program, Champalimaud Center for the Unknown, Lisbon, Portugal; Gatsby Computational Neuroscience Unit. UCL, London

## Abstract

The recently developed ‘two-step’ behavioural task promises to differentiate model-based or goal-directed from model-free or habitual reinforcement learning, while generating neurophysiologically-friendly decision datasets with parametric variation of decision variables. These desirable features have prompted widespread adoption of the task. However, the signatures of model-based control can be elusive – here, we investigate model-free learning methods that, depending on the analysis strategy, can masquerade as being model-based. We first show that unadorned model-free reinforcement learning can induce correlations between action values at the start of the trial and the subsequent trial events in such a way that analysis based on comparing successive trials can lead to erroneous conclusions. We also suggest a correction to the analysis that can alleviate this problem. We then consider model-free reinforcement learning strategies based on different state representations from those envisioned by the experimenter, which generate behaviour that appears model-based under these, and also more sophisticated, analyses. The existence of such strategies is of particular relevance to the design and interpretation of animal studies using the two-step task, as extended training and a sharp contrast between good and bad options are likely to promote their use.

**Author Summary:** Planning is the use of a predictive model of the consequences of actions to guide decision making. Planning plays a critical role in human behaviour but isolating its contribution is challenging because it is complemented by control systems which learn values of actions directly from the history of reinforcement, resulting in automatized mappings from states to actions often termed habits. Our study examined a recently developed behavioural task which uses choices in a multi-step decision tree to differentiate planning from value-based control. Using simulation, we demonstrated the existence of strategies which produce behaviour that resembles planning but in fact arises as a fixed mapping from particular sorts of states to actions. These results show that when a planning problem is faced repeatedly, sophisticated automatization strategies may be developed which identify that there are in fact a limited number of relevant states of the world each with an appropriate fixed or habitual response. Understanding such strategies is important for the design and interpretation of tasks which aim to isolate the contribution of planning to behaviour. Such strategies are also of independent scientific interest as they may contribute to automatization of behaviour in complex environments.

## Introductions

Humans and other animals are thought to use a mixture of different strategies to learn to choose actions that lead to positive outcomes and prevent negative outcomes [1,2]. Much interest is currently focused on the distinction between control systems which employ model-based and (value-based) model-free reinforcement learning (RL) [3–13]. Model-based RL works by learning a predictive model of the specific consequences of actions, and planning by using this model to evaluate the different options prospectively. By contrast, model-free RL directly learns the value of actions through prediction errors, which quantify the difference in worth between actual and expected outcomes. These different strategies offer distinct advantages and disadvantages. Model-based RL is computationally costly, because of the demands of planning many steps into the future. However, it can, in principle, use information efficiently, particularly in the face of a changing environment. This is because the implications that a change has for control in other parts of the environment can be evaluated immediately using the model without having to be directly experienced. Model-free RL incurs little computational cost and supports rapid action selection. However, it is statistically inefficient as it discards information about the specific outcomes of actions, and learns by propagating initially incorrect predictions from states to their sequential predecessors.

Dissociating the contributions of model-based and model-free RL to behaviour is challenging because under many circumstances, including most laboratory based reward guided decision making tasks, they are expected to produce similar behaviour. Outcome devaluation (or, more generally, revaluation) has traditionally been used as a gold-standard test to demonstrate the use of a simple forward model predicting the specific outcomes of actions [1,14,15]. In an outcome devaluation experiment, the subject is trained to perform an action to obtain a reward, e.g. to press a lever for food pellets. The outcome is then devalued in another context, for example by pairing it with illness, and the impact of this devaluation on the subject’s tendency to perform the action is evaluated in extinction, i.e. without further rewards being delivered. If performance of the action is mediated by a model which predicts this specific outcome, devaluing the outcome will reduce the tendency for the action to be performed. If, instead, the subject has simply learnt using model-free reinforcement learning that the action is valuable, without knowing which outcomes underpin that value, devaluation will have no effect on performance in extinction. Research using outcome devaluation paradigms has established that learnt actions are initially specified by model-based RL, but can transition to being devaluation insensitive given extensive training under appropriate conditions [16,17]. This has been interpreted as a shift to model-free RL [4]. Brain regions have been identified whose lesioning or inactivation render animals devaluation insensitive in conditions where intact animals devalue, while a distinct set of regions has been identified as necessary for devaluation insensitive or habitual behaviour [18–29]. This has suggested that model-based and model-free RL are implemented by partially separate neural circuits, and furthermore that learning of each occurs in parallel from the outset of training.

Recent approaches to behavioural neuroscience derive substantial explanatory value from parametric variation of decision variables in the context of large decision datasets. It is therefore desirable to develop tasks which achieve these ends, but also exhibit the critical feature of outcome devaluation – namely the wherewithal to discriminate model-based and model-free RL. The two-step task [6] represents one recently popular approach to creating such a task, attracting a substantial number of human studies [7,10,11,30–40], and with several groups currently adapting the paradigm for use with animal subjects.

At the core of the two-step task is the choice between two actions at the first step. This first action leads to one of two states at the second step; whence reward may subsequently be provided. Each of the actions at the first step has its own preferred state at the second step; however, the transition is probabilistic, such that it occasionally leads to the state preferred by the other action. Depending on whether reward is then delivered, these rare transition trials can provide a form of outcome revaluation, suggesting that model-based and model-free RL might be differentiated according to the different recommendations they give for the ensuing choices. In this paper, we explore the soundness of this differentiation by using simulations to investigate parts of the space of possible strategies that subjects may use to solve the task. In particular, we address two distinct issues which could cause behaviour on the task to be mistakenly identified as arising from model-based RL.

The first issue concerns the stay probability analysis typically used as a tool to assess the relative contributions of model-based and model-free RL. This aims to discriminate these strategies by evaluating the consequences of the transition and outcome on one trial for the frequency with which the choice that led to these events is repeated on the next trial. As the strategies have different update rules through which one trial’s events modify the action values that guide the next trial’s choice, different combinations of trial outcome (rewarded or not) and transition (common or rare) are reinforcing, i.e. increase the chance of the subject repeating the same choice on the subsequent trial. We show that correlation between action values at the start of trials and the subsequent trial events can cause the stay probability analysis, when applied to the behaviour of purely model-free agents, to exhibit a signature classically interpreted as indicative of model-based RL. We further show how the analysis can be modified to correct for these correlations.

The second, and more pernicious, issue arises from the possibility that the state space used by subjects to solve the task may differ from that envisaged by the experimenter. We explore the behaviour of two agents which use extended state representations to solve the task. Again, these may masquerade as being model-based, even though lacking prospective evaluation of the outcomes of actions. The representation employed by the first agent allows it to exploit the predictive relationship that exists between the state from which reward is obtained on one trial and the action that is likely to lead to reward on the subsequent trial. Using model-free RL, the agent learns a fixed mapping from one trial’s events to the next trial’s choice that leads to behaviour that would be assessed as being model-based by either classical or improved stay probability analysis. The representation underlying the second agent makes explicit the latent or hidden state of the world – i.e. which second-step state has higher reward probability. The agent infers this hidden state by observing where it obtains rewards, and uses a fixed mapping from its estimate of the latent state to action. This agent again produces behaviour which is qualitatively very similar to that of a model-based agent. Both agents outperform classical model-free strategies in terms of the fraction of rewarded trials; this provides an incentive for the acquisition of these alternative representations via the ample statistical evidence available particularly to over-trained animal subjects of the correlations that underpin them.

## Results

The results presented in this paper used a reduced version of the two-step task with the state diagram shown in figure 1A. The agent started each trial in the choice state and made a decision between two actions, termed action A and action B. After making this choice the agent reached one of two second-step states termed state *a* and state *b*. Action A normally led to state *a*, and action B normally led to state *b*; however, on a randomly selected 20% of trials, a rare transition occurred, such that action A led to state *b* and action B to state *a*. Unlike in the original two-step task, the agent had no further decision to make in the second step states, but rather had a single action available in each state which led probabilistically to an outcome of reward or no-reward. After receiving either outcome, the agent returned to the choice state to start the next trial. We use a single action in each second stage state both because it is more straightforward to explain the issues we address in this simplified variant and because this version is being used in the current set of rodent variants of the two-step task. The issues we raise are also relevant to task versions with a choice at the second step, though as detailed below they are affected by the reward probability distributions in the second step states.

**Figure 1.**
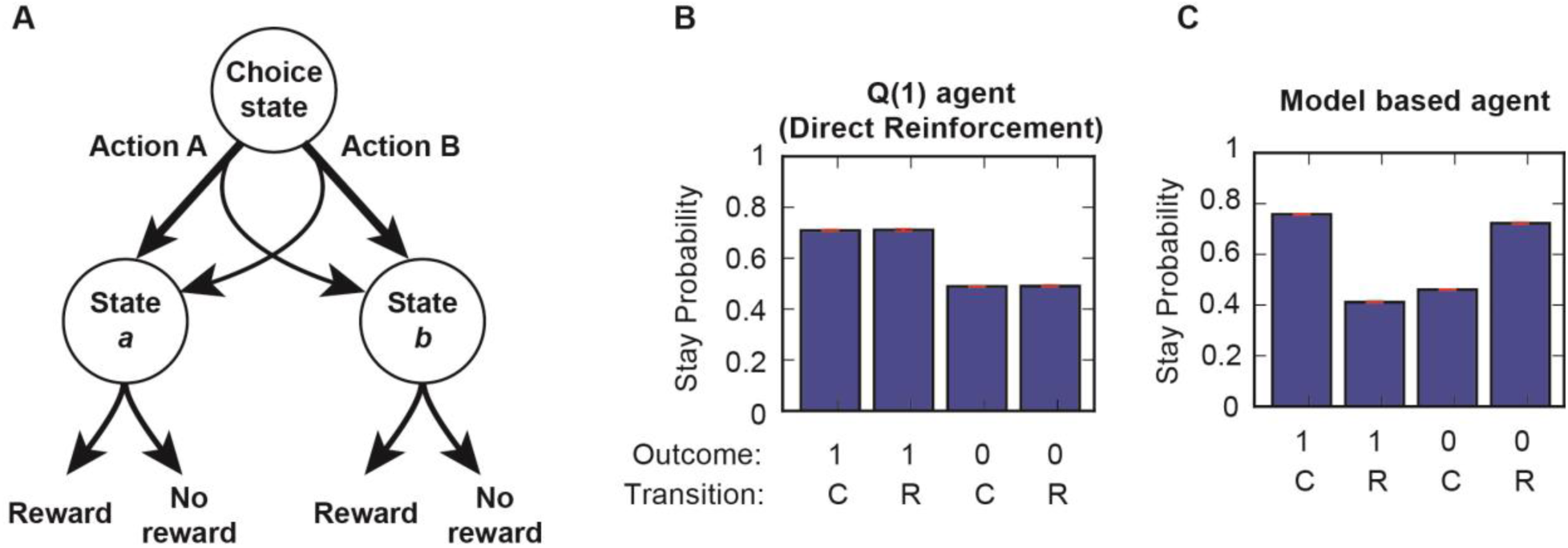
Task diagram and behaviour with neutral reward probabilities. (**A**) Diagram of task state space. (**B, C**) Stay probability plots for *Q*(1)(**B**) and model-based (**C**) agents on a version of the task with fixed equal reward probabilities in states *a* and *b*. Plots show the fraction of trials on which the agent repeated its choice following rewarded and non-rewarded trials with common and rare transitions (SEM error bars shown in red).

We initially simulated the behaviour of a model-free and a model-based agent on a version of the task in which the reward probabilities following both states *a* and *b* were fixed at 0.5. The model-free agent (strictly speaking a *Q*(1)agent, termed a Direct Reinforcement agent in [6]) updated the value of the chosen action (A or B) based on the prediction error between the trial outcome and its current estimate of the action value, using a fixed learning rate. This agent therefore only used information about whether or not the trial was rewarded, and did not use information either about whether a common or rare transition occurred or about the second-step state in which the outcome was received. By contrast, the model-based agent calculated the value of each action A or B as the weighted sum of the values of states *a* and *b*, where the weights were determined by the (known) conditional probabilities of reaching those states after choosing that action. As in [6], model-based and model-free calculations of the values of states *a* and *b* coincide; here we updated the value of the experienced state according to the prediction error between the actual outcome and the previous estimate of that value, using a fixed learning rate.

As reported previously for the two-step task [6], the behaviour of *Q*(1) and model-based agents could be differentiated by looking at how the transition (common or rare) and outcome (rewarded or not) influenced what is called the stay probability, which is the frequency with which the agent repeated the same action on the subsequent trial (Fig. 1B,C). As the action value update used by the *Q*(1) agent is only sensitive to the outcome and not the transition, the stay probability was higher for rewarded than non-rewarded trials, but was not influenced by whether a common or rare transition occurred (Fig. 1B). For the model-based agent (Fig. 1C), a reward following a rare transition increases the value of the state that is more commonly reached from the action that was *not* chosen on that trial. This increases the probability that the agent switches its choice on the subsequent trial. Stay probabilities for the model-based agent therefore showed an interaction between outcome and transition, such that rewards increased stay probability (i.e., were reinforcing) after common transitions, but reduced stay probability after rare transitions.

### Action values at trial start affect stay probabilities

We next simulated the behaviour of the *Q*(1) agent on a version of the task in which the chance of reward reversed every 50 trials between blocks with reward probabilities of 0.8/0.2 in states *a*/*b* and blocks with reward probabilities of 0.2/0.8 in *a*/*b*. The current collection of rodent experiments involves contingencies of this type, rather than the restless bandits of the original human-oriented task [6]. This change in the reward probabilities produced a striking change in the stay probability plot (Fig 2A; repeated for convenient comparison in Fig 3A). Stay probabilities now showed a clear interaction between transition and outcome, though the agent was identical to that used in figure 1B. A logistic regression analysis predicting stay probability as a function of outcome, transition, and transition-outcome interaction confirmed that transition-outcome interaction predicted stay probability (Fig. 2B). This result is counter-intuitive because by construction, the action values and hence choice probabilities of the *Q*(1) agent are unaffected by whether a common or rare transition occurred. The difference in stay probability between trials with the same outcome but different transitions therefore cannot be accounted for by a difference in the action value update that occurred on that trial, as the update is identical irrespective of the transition. Instead, the reason why the action values of the chosen and non-chosen option are (on average) different following trials with the same outcome but different transitions must be that the action values at the start of the trial are (on average) different. This can indeed be seen (Fig. 2C); the mean difference between the action values for the chosen and not chosen option at the start of the trial was larger for common-rewarded and rare-not rewarded trials than for common-not rewarded and rare-rewarded trials.

**Figure 2.**
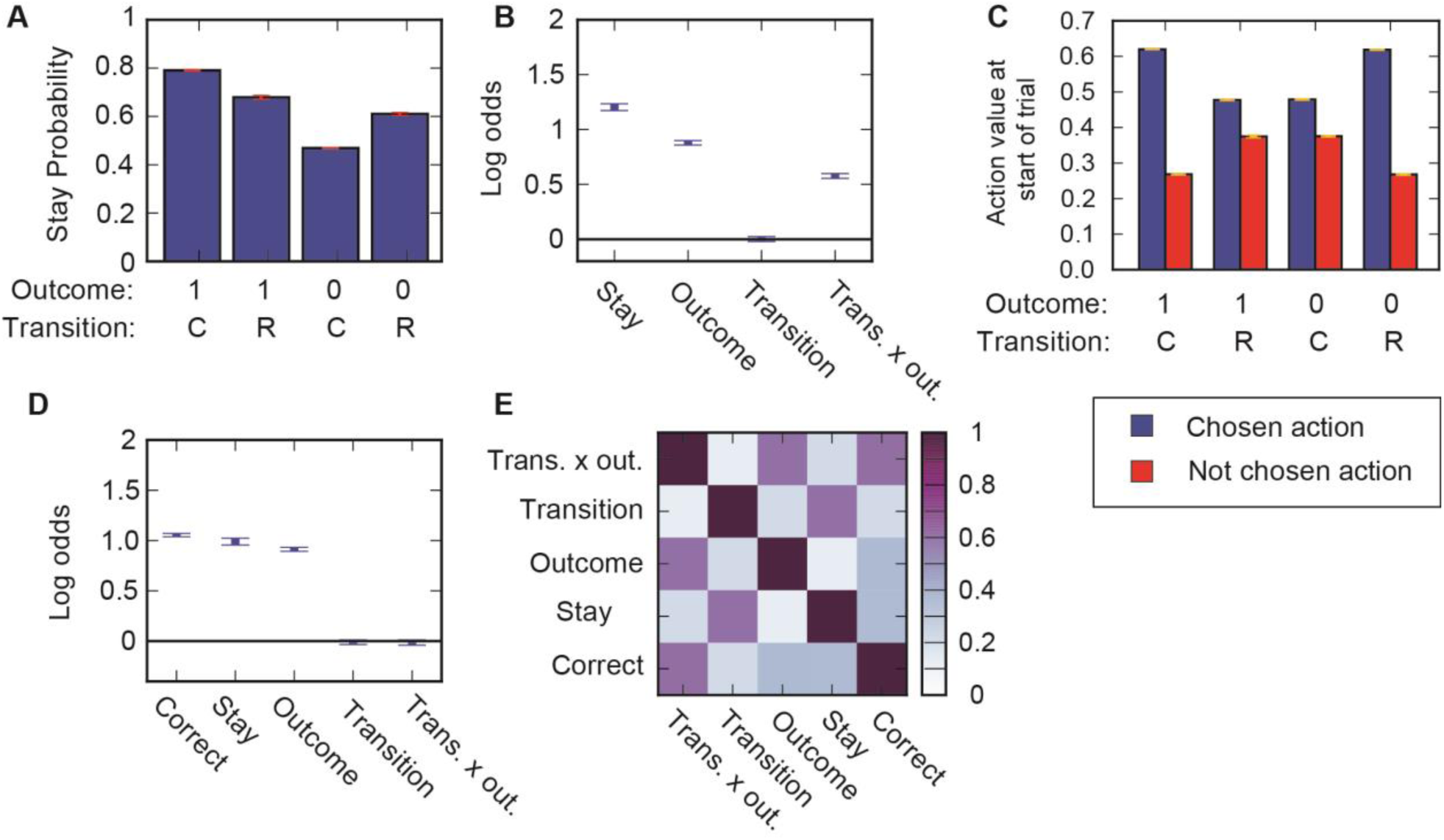
Stay probability transition-outcome interaction for Q(1) agent due to trial start action values. (**A**) Stay probability plot for *Q*(1) agent simulated on version of task with reversals in reward probability for the two second step states. (**B**) Predictor loadings for logistic regression model predicting whether the agent will repeat the same choice as a function of 4 predictors; Stay – a tendency to repeat the same choice irrespective of trial events, Outcome – a tendency to repeat the same choice following a rewarded trial, Transition - a tendency to repeat the same choice following common transitions, Transition x outcome interaction – a tendency to repeat the same choice dependent on the interaction between transition (common/rare) and outcome (rewarded/not). (**C**) Action values at the start of the trial for the chosen and not chosen action shown separately for trials with different transitions (common or rare) and outcomes (rewarded or not). Yellow error bars show SEM across sessions. (**D**) Predictor loading for logistic regression model with additional predictor capturing tendency to choose the action which was correct on the previous trial, i.e. the action whose common transition lead to the state which currently has high reward probability. (**E**) Across trial correlation between predictors in logistic regression analysis shown in (**D**).

**Figure 3.**
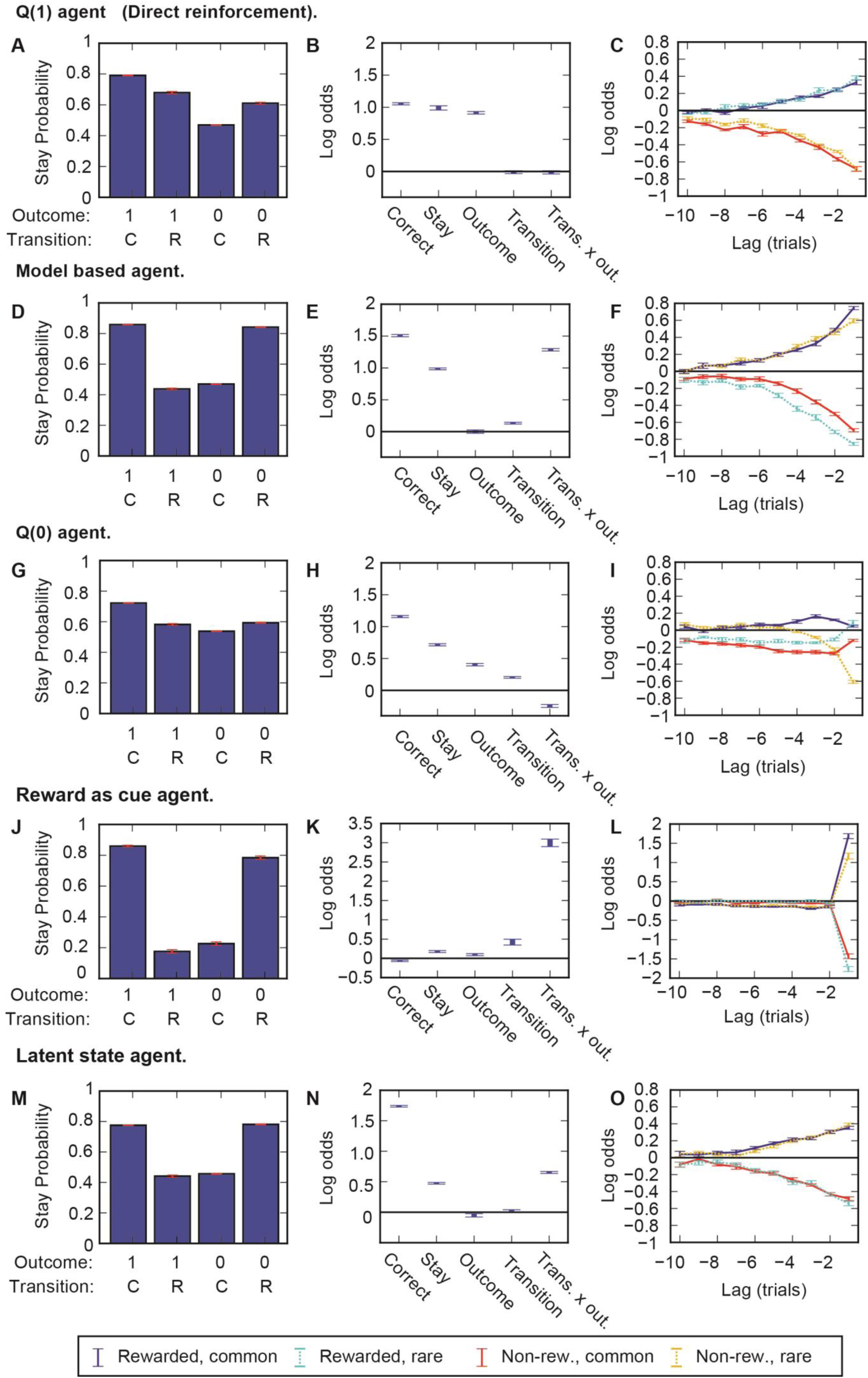
Comparison of agents’ behaviour. Comparison of the behaviour of all agents types discussed in the paper on version of task with 0.8/0.2 reward probabilities with reversals every 50 trials. Left panels – Stay probability plots. Centre panels - Predictor loadings for logistic regression model predicting whether the agent will repeat the same choice as a function of 5 predictors; Correct - a tendency to choose the action which was correct on the previous trial, Stay – a tendency to repeat the same choice irrespective of trial events, Outcome – a tendency to repeat the same choice following a rewarded trial, Transition - a tendency to repeat the same choice following common transitions, Transition x outcome interaction – a tendency to repeat the same choice dependent on the interaction between transition (common/rare) and outcome (rewarded/not). Right panels - Predictor loadings for lagged logistic regression model. The model uses a set of 4 predictors at each lag, each of which captures how a given combination of transition (common/rare) and outcome (rewarded/not) predicts whether the agent will repeat the choice a given number of trials in the future, e.g, the ‘rewarded, rare’ predictor at lag -2 captures the extent to which receiving a reward following a rare transition predicts that the agent will choose the same action two trials later. Legend for right panels is at bottom of figure. Error bars in all plots show SEM across sessions.

Why are action values at the start of the trial correlated with subsequent trial events, specifically the transition-outcome interaction? There are two steps in the argument. First, the agent is more likely to choose the correct action, i.e. the action which commonly leads to the state with high reward probability, on trials where there is a large difference in action values between the two options. On trials in which the difference in action values is small, the agent has little evidence that one option is better than the other, and has a higher change of choosing the incorrect action. Second, choosing the correct, rather than incorrect, action changes the probabilities of observing different combinations of trial events. Rewarded common transitions and unrewarded rare transitions are more likely to occur following a correct action than they are to occur following an incorrect action. Conversely, rewarded rare transitions and unrewarded common transitions are more likely to occur following an incorrect action.

To summarise; the difference in action values going into the trial correlates with the probability of choosing the correct option. Whether the agent chooses the correct option determines the probability of observing each combination of subsequent trial events. Therefore when trials are divided into groups by outcome and transition, the action values at the start of the trial show a transition-outcome interaction (Fig. 2C), which is then also observed for the stay probabilities (Fig. 2A), even though the agent did not use any information about the transition in its action value update. This effect is not restricted to block based reward probabilities; it can also be observed when reward probabilities change as random walks (Fig. S1A) or with static fixed reward probabilities of 0.8 / 0.2 in states *a* / *b* (Fig. S1B), rather than 0.5/0.5 as in Fig. 1.

It is possible to modify the logistic regression analysis of stay probabilities to prevent differences in action values at the start of the trial from appearing as a spurious loading on the transition-outcome interaction predictor. This can be done by including an additional ‘correct’ predictor which captures the tendency of the agent to choose the option which was the correct choice on the previous trial. Including this additional predictor completely removed loading on the transition-outcome interaction predictor for the *Q*(1) agent (Fig. 2D; repeated for convenient comparison in Fig 3B), correctly revealing that only the trial outcome affected the agent’s subsequent choice. For a model-based agent simulated on the same task version (with reversals in reward probabilities), this extended logistic regression analysis showed positive loading on the transition-outcome interaction predictor (Fig. 3E) reflecting the true importance of this interaction to the action value update used by the agent. The addition of a ‘correct’ predictor works because the correlation between actions values at the start of a trial and the subsequent transition-outcome interaction is entirely mediated by the correlation between these action values and whether the agent chose the correct action on that trial. Explicitly including a predictor for choosing the option which was the correct choice on the previous trial absorbs the variance due to action values going into the trial which would otherwise be absorbed by the transition-outcome interaction predictor due to correlation between these two predictors (Fig. 2E).

An alternative way of differentiating model-based and model-free strategies is a lagged logistic regression analysis which examines the effect on choice probability of trial events at different lags relative to the current trial (Miller at al. Soc. Neurosci. Abstracts 2013, 855.13). Figures 3C,F show a lagged logistic regression analysis for *Q*(1) and model-based agents. The analysis evaluated how different combinations of outcome and transition predict that the agent will repeat the same choice a given number of trials in the future. For example, the ‘rewarded, rare’ predictor at lag -2 captures the extent to which receiving a reward following a rare transition predicted that the agent will choose the same action two trials later. This analysis is therefore an extension of the classical stay probability analysis to include the effect of earlier trials. For the *Q*(1) agent (Fig. 3C), obtaining a reward predicted that the agent will repeat the same choice irrespective of the transition, with a smoothly decreasing predictive weight at increasing lag. For the model-based agent (Fig. 3F), rewarded-common transitions and non-rewarded rare transitions predicted the agent will repeat the same choice, while rewarded-rare and non-rewarded common transitions predict the agent will not repeat the same choice, again with the predictive weight smoothly decreasing with increasing lag.

Although a *Q*(1) agent is typically used to illustrate model-free behaviour on the two-step task, it represents one end of a spectrum of model-free agents differentiated by the extent to which the action value update at the first step depends on either the trial outcome or second step action values. This spectrum is parameterized by the eligibility trace parameter conventionally called λ. The update used by the *Q*(1) agent depends only on the trial outcome and not at all on the values of the second step state (or second-step actions in the original two-step task [6]). At the other end of the spectrum is the *Q*(0)agent which updates the value of the first step action based only on the value of the second-step state, with no direct influence of the trial outcome. The value of the second-step state is then updated based on the trial outcome. The behaviour of a *Q*(0) agent on the simplified two-step task is shown in figure 3G-I, and the behaviour of model-free agents which use mixtures of the *Q*(1) and *Q*(0) updates are shown in figure S1. The one trial back logistic regression analysis for the *Q*(0) agent (Fig. 3H) shows positive loading on the transition and outcome predictors and negative loading on the transition-outcome interaction predictor. Loading on the transition-outcome interaction predictor in the extended logistic regression analysis distinguishes the model-based agent from all of the model-free agents, none of which shows positive loading on this predictor. The lagged logistic regression for the *Q*(0) agent shows a complex pattern in which the predictive weight of each combination of trial events does not decay smoothly at increasing lags.

### Extended state representations

We have so far considered how action values at the start of the trial can affect the stay probability based analyses used to differentiate model-based from model-free behaviour on the two-step task. We now turn to a different source of ambiguity about behaviour on the task: the existence of strategies which use state representations that differ from the conventional set of states used to define the task structure. It turns out that model-free versions of these can also produce behaviour similar to that of a model-based agent without using the prospective action evaluation that is the hallmark of classical model-based RL.

The two-step task has a circular structure in which subjects cycle repeatedly through the decision state, second step states and trial outcomes. This repeating structure provides opportunities for subjects to learn predictive relationships between events on one trial and the actions that are likely to lead to reward on the subsequent trial. One such predictive relationship is that the location where reward is obtained on one trial predicts which choice on the next trial is likely to lead to reward. That is, if a reward is obtained in state *a*, the reward probability is higher for choosing action A on the subsequent trial, while if reward is obtained in state *b* the reward probability is higher for choosing action B on the subsequent trial. Note that this predictive relationship holds true across reversals in which second step state has higher reward probability. The locations where reward is obtained, and conversely where non-rewards are obtained, can therefore, in principle, be used as discriminative stimuli to guide choice on the next trial. We therefore considered the behaviour of a ‘reward-as-cue’ agent which uses the location of reward as a discriminative stimulus for the state of the world. Specifically, the reward-as-cue agent treated the choice between actions A and B as occurring in one of 4 distinct states on each trial, defined by whether a reward or non-reward was obtained in state *a* or *b* at the end of the previous trial. The agent used model-free RL to learn independent values of actions A and B in each of these 4 states. Like the *Q*(1) agent, the reward-as-cue agent updated the value of the chosen action dependent on reward prediction error between its current estimate of the action value and the trial outcome, without using the action value at the second step in the update. The agent learned action values which produced the strategy of choosing action A following rewards in state *a*, action B following rewards in state *b*, action B following no reward in state *a* and action A following no reward in state *b*. This corresponds to a strong stay probability transition-outcome interaction (Fig. 3.J,K). Unlike the other agents considered so far, the reward-as-cue agent does not adapt to changes in the reward probabilities across blocks through changes in its action values. Rather, the action values are stable across blocks and reflect a fixed mapping between where reward is obtained and which action should be taken on the next trial.

It does not seem unreasonable to assume that over-trained animals could learn to use the location of reward as a discriminative stimulus to guide choice on the next trial, as animals straightforwardly learn to use discriminative sensory stimuli of various sorts as cues for the best action to take next [41–44]. Once learnt, this strategy would be minimally cognitively demanding as it is essentially a fixed stimulus-response habit with only a limited demand on working memory. However, although the reward-as-cue agent gives behaviour on the one trial back stay probability analysis which is qualitatively similar to that of a model-based agent, it shows a very different pattern of loadings in the lagged logistic regression analysis (Fig. 3L). Rather than the smooth drop off of predictive weight with increasing lag observed for the model-based agent, only the previous trial events are predictive of the reward-as-cue agent’s behaviour.

**Figure 4.**
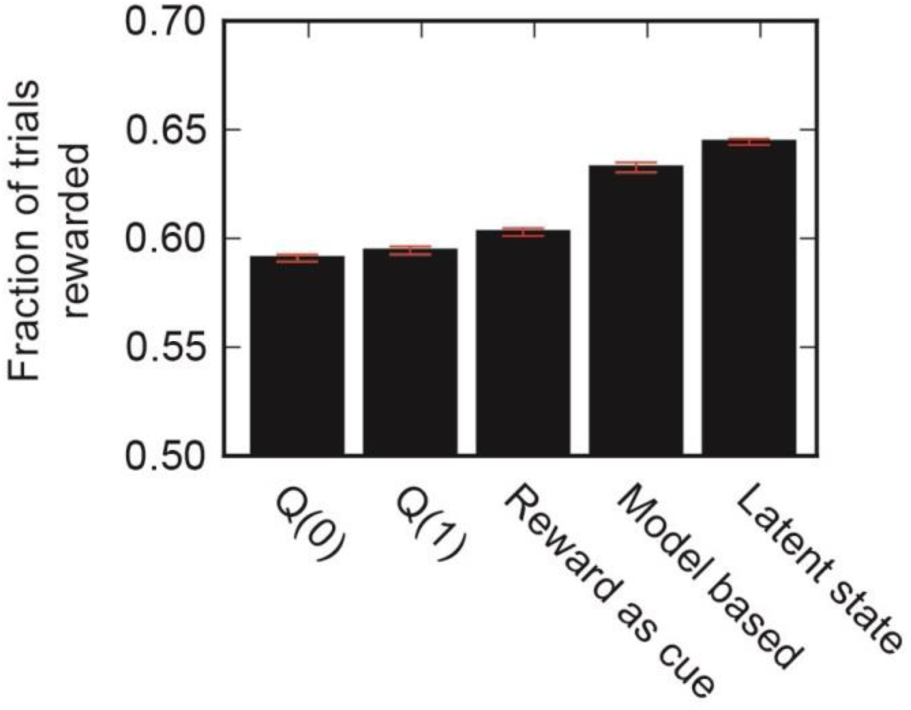
Comparison of agents’ performance. Performance achieved by different agent types with parameter values optimised to maximise the fraction of trials rewarded. For the reward a cue agent, performance is shown for a fixed strategy of choosing action A (B) following reward in state *a* (*b*) and action B (A) following non-reward in state *a* (*b*). SEM error bars shown in red.

For all the agents in Fig. 3, we chose parameters determined by a maximum likelihood fit to the behaviour of the model-based agent, so that they would all have comparable average behaviour; see materials and methods). For the reward-as-cue agent, this suggested a very low learning rate (0.002). If, instead, we chose parameters for all agents that maximized the fraction of trials that are rewarded, the reward-as-cue agent outperformed both *Q*(1) and *Q*(0) model-free agents (Fig. 4).

The reward-as-cue strategy works because there is in fact a latent, unobservable state of the world that is important to the decision problem – whether the reward probability is higher in state *a* or *b*. The location where reward is obtained is correlated with, and hence informative about, this latent state, and therefore can be utilised as a discriminative stimulus to guide behaviour. However, because the reward-as-cue strategy uses only the most recent reward as a discriminative stimulus, it is far from optimal. We therefore evaluated the behaviour of a different agent we term ‘latent-state’ which understands that the world is always in one of two latent states, one in which the reward probability is high in state *a* and low in state *b*, and the other in which the reward probability is high in state *b* and low in state *a*. At the end of each trial the latent-state agent performed a Bayesian update of its estimate of the probabilities that world is in each latent state based on the observed trial events. In updating the probabilities the agent also assumed that there is a small probability (the inverse of the mean block length) that the state of the world switches between trials. This amounts to the assumption that the block lengths are exponentially distributed, rather than being of fixed length, as generally employed. We did not explicitly model the learning of action values in each of these latent states, but rather assumed asymptotic behaviour in which the agent chose action A with high probability in the latent state where state *a* had high reward probability and action B with high probability in the latent state where state *b* had high reward probability.

The behaviour of the latent-state agent looked qualitatively very similar to that of the model-based agent. The one trial back stay probability analyses showed a transition-outcome interaction (Fig. 3M-N). As for the model-based agent (Fig. 3F), the lagged logistic regression analysis for the latent-state agent (Fig. 3O) showed a tendency to repeat choices that were followed by rewarded common and non-rewarded rare transitions, and to not repeat choices that were followed by non-rewarded common and rewarded rare transitions, with a gradually decreasing predictive weight at increasing lag. However, the behaviour of the latent-state and model-based agents could be discriminated using model fitting, with data simulated by the model-based agent being fit with higher likelihood by the model-based agent (Fig. 5A) and data simulated by the latent-state agent being fit with higher likelihood by the latent-state agent (Fig. 5B). Data simulated from either latent-state or model-based agents was better fit by both of these agents than by either the *Q*(0) or *Q*(1) model-free agents. With parameters optimized to maximize reward, the latent-state agent, embodying as it does a faithful characterization of the environment, achieved the highest performance in terms of fraction of trials rewarded of all the agent types examined (Fig. 4).

**Figure 5.**
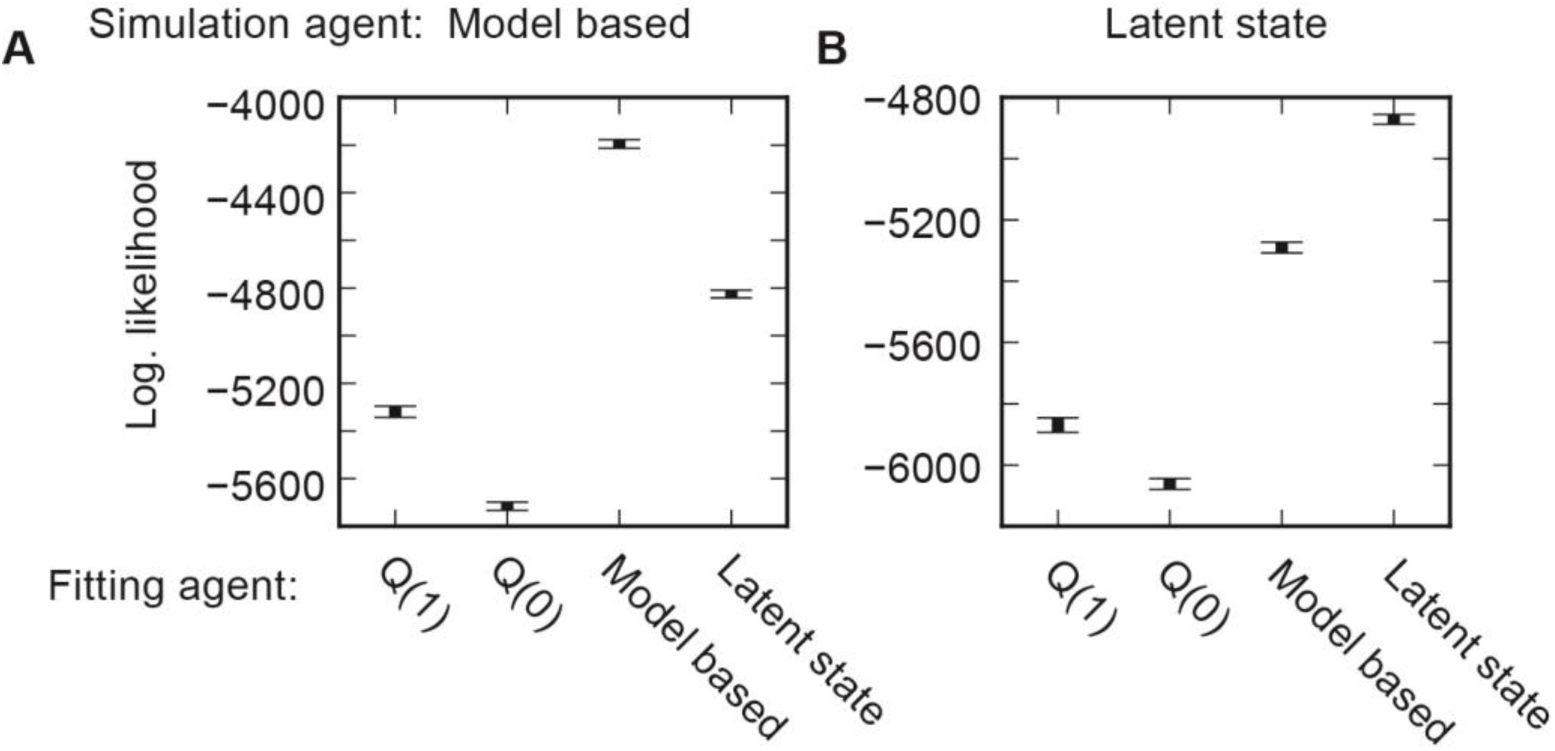
Likelihood comparison. Data likelihood for maximum likelihood fits of different agent types to data simulated from a model-based agent (**A**) or a latent-state agent (**B**).

## Discussion

We have identified two ways in which behaviour on the two-step task could be incorrectly identified as arising from prospective model-based evaluation of actions. The first issue is with the stay probability analysis commonly used as a metric of subjects’ strategies. We showed that rather than reflecting only the action value update occurring on a given trial, which is distinct for model-based and model-free action evaluation, stay probabilities are also affected by action values at the start of the trial. This can cause the behaviour of a model-free agent to exhibit a stay probability transition-outcome interaction, which has previously been interpreted as a signature of model-based behaviour. The second issue is the existence of alternative strategies which use different state representations from the basic states that define the task structure and produce behaviour which is similar to that of a model-based agent though not dependent on prospective evaluation of the outcome of actions.

Despite a substantial body of literature employing the two-step task [6,7,10,11,30–40], the influence of trial start action values on stay probabilities has not been previously recognised. This is because the quantitative details of the task design strongly influence the strength of this effect, primarily through their influence on the magnitude of the average difference between reward probabilities in the two second step states (*a*, *b*). When this difference is zero, stay probabilities for the *Q*(1) agent depend only on trial outcome (Fig.1B). In the original version of the task [6], each second step state afforded two actions, whose reward probabilities varied according to reflecting Gaussian random walks on the interval [0.25, 0.75]. The difference on average between the reward probabilities of the better action in each second step state was therefore substantially smaller than that shown in the simulations here. The effect size also depends on the probability of making a common transition, being smaller when the common transition probability is closer to 0.5. In the original task this probability was 0.7, while in the version used in these simulations it was 0.8. In figure S3 we show the stay probability analysis for a *Q*(1) and a model-based agent with the same parameters used in figure 3, simulated on the version of the task used in [6]. The *Q*(1) agent shows a very small transition-outcome interaction while the model-based agent shows a strong interaction. The influence of trial start action values on stay probabilities is therefore unlikely to be an issue in interpreting stay probability plots in published work using the original version of the task. However, the conditions under which the effect size is negligible; common transition probabilities close to 0.5 and small differences in reward probabilities between the two second step states, are also those conditions under which is impossible to do much better than chance by making optimal choices at the first step. In our experience training animals, if the differences between good and bad options are small, subjects will often switch to behavioural strategies such as always choosing the same option, or alternation, which obtain rewards at chance level with less cognitive effort than even the simplest of the strategies we have analysed. If the two-step task is adapted for use with animals by increasing the contrast between good and bad options, the classical stay probability analysis ceases to be a viable tool for assessing how events on one trial affect choice on the subsequent trial due to the influence of trial start action values.

We demonstrated that the effect of trial start action values can be corrected for in a logistic regression analysis of stay probabilities by including a predictor which captures the tendency to choose the action which was correct on the previous trial, i.e. to repeat correct choices. We suggest this modified stay probability analysis will be a useful tool for evaluating the influence of trial events on subsequent choice in task variants where the classical stay probability analysis gives misleading results. One cost of using this additional predictor is that, as it is correlated with the transition-outcome predictor (Fig. 2E), the relative loadings on these two predictors will be sensitive to fluctuations in the data, potentially requiring larger datasets to achieve reliable results. The addition of a binary ‘correct’ predictor also does not completely absorb the influence of trial start action values on stay probabilities in situations where the reward probabilities change as random walks rather than reversals between two block types, presumably because it does not capture the magnitude of the difference in reward probabilities between the two options. With random walk reward probabilities, using the continuous valued difference in reward probabilities works better than a binary ‘correct’ predictor to compensate for trial start action values (data not shown).

The second issue we have identified with the two-step task is that, due to its repeating structure, subjects could, in principle, learn to exploit correlations between events on a given trial and reward probabilities on future trials to produce behavioural strategies that look similar to model-based behaviour but do not use prospective evaluation of actions. One simple strategy which we termed ‘reward-as-cue’ uses model-free reinforcement learning to exploit the predictive relationship between where rewards are obtained on a given trial and which action on the subsequent trial has higher reward probability. This strategy learns a fixed mapping between events on one trial and choice on the next trial (e.g. reward in state *a* → choose action A). Notably, it outperformed classical model-free strategies in terms of acquiring reward. The attraction of this strategy from the point of view of the subject is that once learned it requires no further updating of action values to adjust to changes in reward probability in the two second-step states, and hence can be fully automatized into a stimulus-response habit. Though this strategy produces a strong stay probability transition-outcome interaction, it can be distinguished from model-based behaviour as only the most recent trial influences choice. However behaviour strikingly similar to that of a model-based agent was generated by a more sophisticated strategy we termed ‘latent-state’, which uses the location of recent rewards as a discriminative stimulus for which of two latent states the world is in (high reward probability in state *a*, or high reward probability in state *b*), and follows a fixed mapping from the latent state of the world to choice. Behaviour simulated from latent-state and model-based agents could be differentiated by model-fitting. However, given the similarity of the behaviour they generate, model fitting may not be a reliable way of discriminating these strategies in the wild where they may be expressed in combination with other strategies and in forms which do not exactly match the quantitative details of the fitted model. If subjects do in fact use latent state strategies, this would substantially complicate using the two-step to study classical model-based control employing prospective action evaluation.

Is it plausible that subjects could learn latent-state type strategies in the two-step task? There is evidence that in probabilistic reversal learning tasks, humans [45] and monkeys [46] learn that there are in fact two distinct latent states of the world and use inference about the current latent state to guide their behaviour. A further reason to think that the latent state inference at the heart of this strategy is plausible is that subjects in tasks that require integration of noisy sensory evidence appear successfully to perform a close analogue. Examples of such tasks include random dot motion discrimination and the Poisson clicks auditory discrimination task [41,43]. Consider the latter: in this task, the subject receives two Poisson like trains of clicks, one presented on the left and one on the right. The subject must integrate information from these trains to judge whether the rate of clicks is higher on the right or the left, and use this information to make a binary choice between two possible actions. The two-step latent state strategy is equivalent, only with integration across, rather than within, trials. That is, the subject (over multiple trials) receives two trains of outcomes, one in each of the two second step states. The subject must integrate these noisy signals over time to evaluate which second step state has higher reward probability, and use this information to guide a binary decision. In both cases there is a latent state of the world which is important for choice – whether the click rate is higher on the left or right in the Poisson clicks, and whether the reward rate is higher in state *a* or *b* in the two-step. Both cases require integration of noisy evidence over time to evaluate this latent state and the use of this information to guide a binary decision. Certainly there are important differences between these tasks; the timescale of integration is longer in the two-step and spans multiple trials, the discriminative stimuli in the two-step are themselves rewards, and subject take an active role in sampling the two information streams. However, there are also striking commonalities and it therefore does not seem implausible that with sufficient training animal and human subjects, both of which readily learn evidence integration tasks, could learn latent state based strategies on the two-step task.

Both the reward-as-cue and latent-state strategies (termed collectively ‘extended-state’ strategies) work by exploiting the regularity in the task structure that the location where rewards are obtained correlates with which first step action has higher reward probability. Evidence for this regularity accrues slowly as it is only across multiple reversals in the reward probabilities that the correlation becomes apparent. It therefore seems probable that if subjects do learn to exploit this regularity, the strategy would only arise after extended experience with the task. In the original version of the task used typically in the human literature, subjects do a total of ∼200 trials. The limited number of trials performed, and the fact that human subjects have been trained to understand the true task structure - presumably priming the use of a model-based strategy - both argue against the possibility that the apparently model-based behaviour reported in the human literature in fact arises from extended-state strategies. However, in the many adaptations of the task to animal subjects, the existence of alternative strategies is more problematic. Animal subject are often extensively trained on tasks before recordings or manipulations are performed, providing ample opportunity to learn task regularities. Additionally, as discussed earlier, to ensure that animal subjects track changes in reward probabilities the contrast between good and bad options may have to be increased from that in the original task. Higher contrast between reward probabilities provides stronger statistical evidence for the regularity that underpins extended-state strategies. We therefore think that the possibility that subjects may be utilising such strategies should be taken seriously in the design and interpretation of experiments using two-step type tasks.

What options exist to minimise the probability that apparently model-based behaviour is in fact due to such strategies? One option would be to avoid overtraining subjects, limiting the total number of trials they perform. This may be a good approach for some applications, however it is often desirable to generate very large behavioural datasets to better quantify the effect of manipulations or the relationship between behaviour and neural activity. Indeed, a substantial part of the appeal of the two-step task when compared with outcome devaluation is that it apparently offers the ability to dissociate model-based and model-free control during ongoing behaviour rather than a brief single shot test. A second possibility is to modify the reward probabilities to reduce the salience of the correlations underpinning extended-state strategies. As the contrast between good and bad options is reduced, these correlations become weaker, but subjects may disengage with the task and stop tracking reward probabilities. A third possibility is to accept that it may be difficult to disambiguate extended-state from classical model-based strategies purely from behaviour, and use neural data to try and disambiguate the strategy used by subjects. Latent state strategies go beyond classical model-free RL and are interesting in their own right. However, this concession does reduce some of the reductionist appeal of the two-step task as a tool for dissociating behavioural strategies. A fourth possibility would be to utilise tasks which do not have a repeating structure. A number of tasks have recently been developed to study planning in which subjects choose trajectories through multi-state environments to obtain reward, but are not returned to the same state at the start of each trial [9,13,47]. Such tasks do not appear susceptible to the relatively simple extended-state strategies which prosper in the two-step task.

A final option that we think is promising is to modify the two-step task by introducing reversals into the transition matrix which maps the first step choice to the state reached at the second step. In this task variant, not only does the reward probability in each second step state change over time, but the action which must be chosen to reach a given second step state also changes. This modification substantially increases the complexity of latent state strategies while minimally changing the complexity of classical model-based control. Reversals in the transition matrix break the fixed predictive relationship in the original task between where reward is obtained and which action at the first step is likely to lead to reward. To solve this version through a fixed mapping from an inferred latent state to action requires latent states that are non-linear combinations of where rewards have been obtained and which actions have led to which states. Classical model-based control continues to work as normal, with the proviso that the subject must continually learn from experience the current mapping from first-step actions to second-step states.

The possibility we have identified here for model-free strategies to masquerade as model-based mirrors proposals that apparently model-free behaviour on the two-step task may in fact be due to model-based selection applied to action sequences [11]. Though very different in their underlying mechanisms, both indicate the complexity of cleanly dissociating the contribution of different learning strategies to behaviour.

The two-step task latent state strategy provides an example of how agents may turn a planning problem into a set of automatized state-response mappings if there is a limited set of relevant states of the world, each with their own appropriate response. Even if the planning problem is large, with a great diversity of possible solutions, e.g. navigating from home to work, with experience the decision may be automatized to a mapping from a small number of relevant states of the world, e.g. is it rush hour, to a set of options which are known to work best in each condition. Such automatization is more sophisticated than stimulus-response habits as typically envisioned; the states of the world that evoke the response may be high level abstractions rather than directly observable stimuli, and the responses may be action sequences, or options in the hierarchical RL formalism [48]. However, as cached state – action mappings learnt through a history of reinforcement, such strategies have commonalities with classical habits and may be learnt using similar model-free RL algorithms applied to higher level state and action representations, perhaps instantiated in cortical-basal ganglia loops involving higher level cortices and associative and limbic striatal sub-regions. These considerations bring to the forefront the question of what state representations are learned and used [49,50], something known to be central to the speed with which agents learn to solve decision problems. As we have explored here, it is also important for the design and interpretation of behavioural experiments, and is only likely to become more so as increasingly complex questions are asked of behaviour.

## Materials and Methods

### Simulations

All simulations and analysis were conducted in Python. Full code used to produce the paper figures is available at *https://bitbucket.org/takam/two-step-simulations*. For each agent, 10 sessions of length 10000 trials were simulated. Where errorbars are used these show standard error of the mean across session.

### Agents

In describing the action value updates used by the different agents we use the following variables:

*Q*(*a*_1_): The value of the first step action chosen on the trial.

*Q*(*a*_2_): The value of the second step action that was reached on the trial.

*r*: The trial outcome (1 for reward, 0 for non-reward).

*α*: The agent’s learning rate.

All agents except the latent-state agent used a softmax decision rule with inverse temperature parameter *T* to determine choice probabilities as a function of action values.

Action value updates for these agents were as follows:

### Q (λ) agents

The action value update rules used by the *Q*(λ) agents (including *Q*(0) and *Q*(1) agents) were:

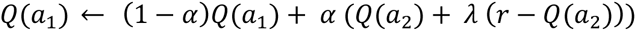

Where λ is the eligibility trace parameter (0 for the *Q*(0) agent and 1 for the *Q*(1) agent).

*Model-based agent:*

The action value update rule used by the model-based agent at the second step was:

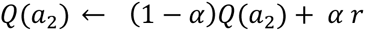

At the start of each trial the agent computed action values for the first step actions as:

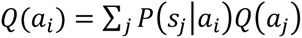

Where *Q*(*a*_*i*_) is the value of first step action *i*, *Q*(*a*_*j*_) is the value of the action available in second step state *j*, and *P*(*s*_*j*_|*a*_*i*_).is the true probability of reaching second step state *j* after choosing action *i*.

### Reward-as-cue agent

The reward-as-cue agent treated the choice between actions A and B as occurring in one of four different states on each trial, corresponding to the 4 combinations of the outcome (1 or 0) and second step state (*a* or *b*) that occurred on the previous trial. The action value update used was:

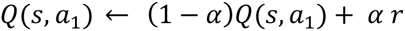

Where *Q*(*s*, *a*_1_) is the value of the action chosen at the first step in the relevant state.

### Latent-state agent

The latent-state agent believed there were two states of the world, one of which had reward probabilities of (0.2, 0.8) in second step states *a* and *b* respectively, and the other with reward probabilities of (0.8, 0.2). At the start of each trial the agent performed a Bayesian update of the probability that the world was in each of these states based on the previous trial events. The agent then updated the probability that the world was in each state to account for the possibility that the world reversed in state between the previous and current trial, which was assumed to occur with probability ω. The agent used a probabilistic mapping from its estimate of the state of the world to choice, choosing with probability (1 – ε) the action with higher reward probability in the most probable state, and with probability ε the action with higher reward probability in the less probable state.

### Parameter values

The parameter values of the model-based agent were set to: *α* = 0.5, *T* = 5

To ensure that average behaviour for the different agents was comparable, the parameters of the other agents were set by maximum likelihood fitting to data simulated from the model-based agent. This resulted in the following agent parameters:

*Q*(0) agent: *α* = 0.506, *T* = 2.90

*Q*(1) agent: *α* = 0.327, *T* = 3.21

*Q*(0.25) agent: *α* = 0.503, *T* = 3.46

*Q*(0.5) agent: *α* = 0.477, *T* = 3.55

*Q*(0.75) agent: *α* = 0.403, *T* 3.41

Reward-as-cue agent: *α* = 0.00229, *T* = 6.52

Latent-state agent: *ω* = 0.0364, *ε* = 0.190

### Comparing agent performance

To evaluate the performance of the different agents in figure 4, agent parameter values were optimised using a grid search to maximise the fraction of trials that were rewarded. For the Reward-as-cue agent we used the performance of a deterministic reward-as-cue strategy which choose option A following reward in state *a*, option B following reward in state *b*, choose option A following non-reward in state *b*, option B following non-reward in state *a*.

### Logistic regression analysis

In all logistic regression analyses, the dependent variable was the subject’s choice, coded as stay vs switch. For the basic one trial back logistic regression, the explanatory variables used were the outcome of the previous trial (rewarded or not), the transition (common or rare) and their interaction. The constant term in the regression capturing the subjects overall tendency to stay was plotted with the other predictors and termed Stay. For the extended one trial back logistic regression analysis a further binary explanatory variable termed ‘Correct’ was added which indicated whether the previous choice was correct, i.e. had the higher probability of leading to reward. For the lagged regression analysis, a set of four binary explanatory variables was used at each lag. These were termed Rewarded common, Rewarded rare, Non-rewarded common and Non-rewarded rare. The Rewarded-common variable indicated whether the previous trial was rewarded and had a common transition and the other predictors indicated respectively whether the other combinations of outcome and transition had occurred.

## Acknowledgements

The authors thank Evan Russek, Kevin Miller, Bruno Miranda, Eric DeWitt, Nathaniel Daw and Anthony Dickinson for useful discussions.

## Supplementary Figures

**Figure S1.**
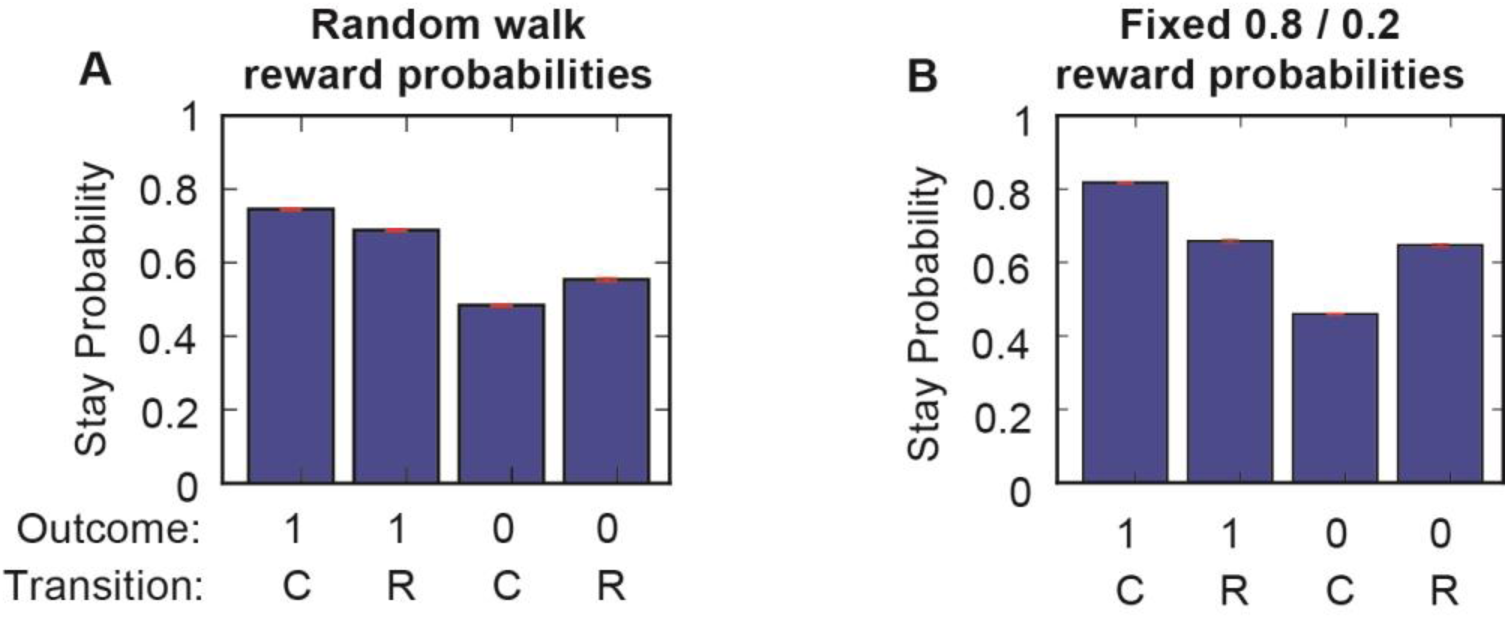
Q (1) agent with random walk and fixed reward probabilities. Stay probability plot for *Q*(1) agent simulated on; (**A**) version of the task where reward probabilities change as reflecting random walks on the range 0 – 1, (**B**) version of the task with fixed reward probabilities of 0.8 / 0.2 in states *a* / *b*.

**Figure S2.**
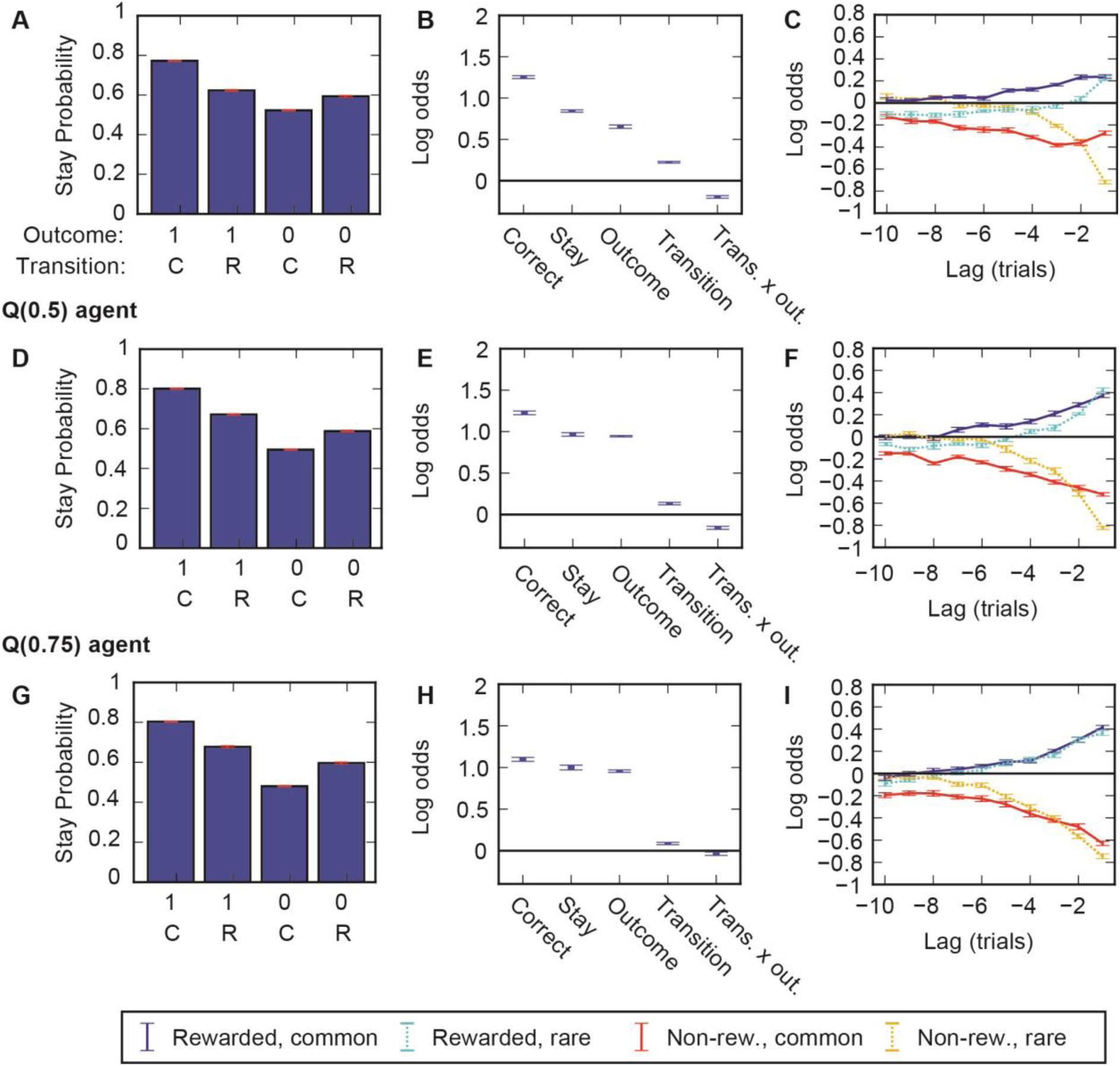
Intermediate values of lambda. Comparison of behaviour from agents with intermediate values of the lambda parameter that controls the relative contribution of the *Q*(1) update and TD0 update. Left panels - stay probability plots. Centre panels - One trial back logistic regression, Right panels - lagged logistic regression. Legend for right panels is at bottom of figure. Error bars in all plots show SEM across sessions.

**Figure S3.**
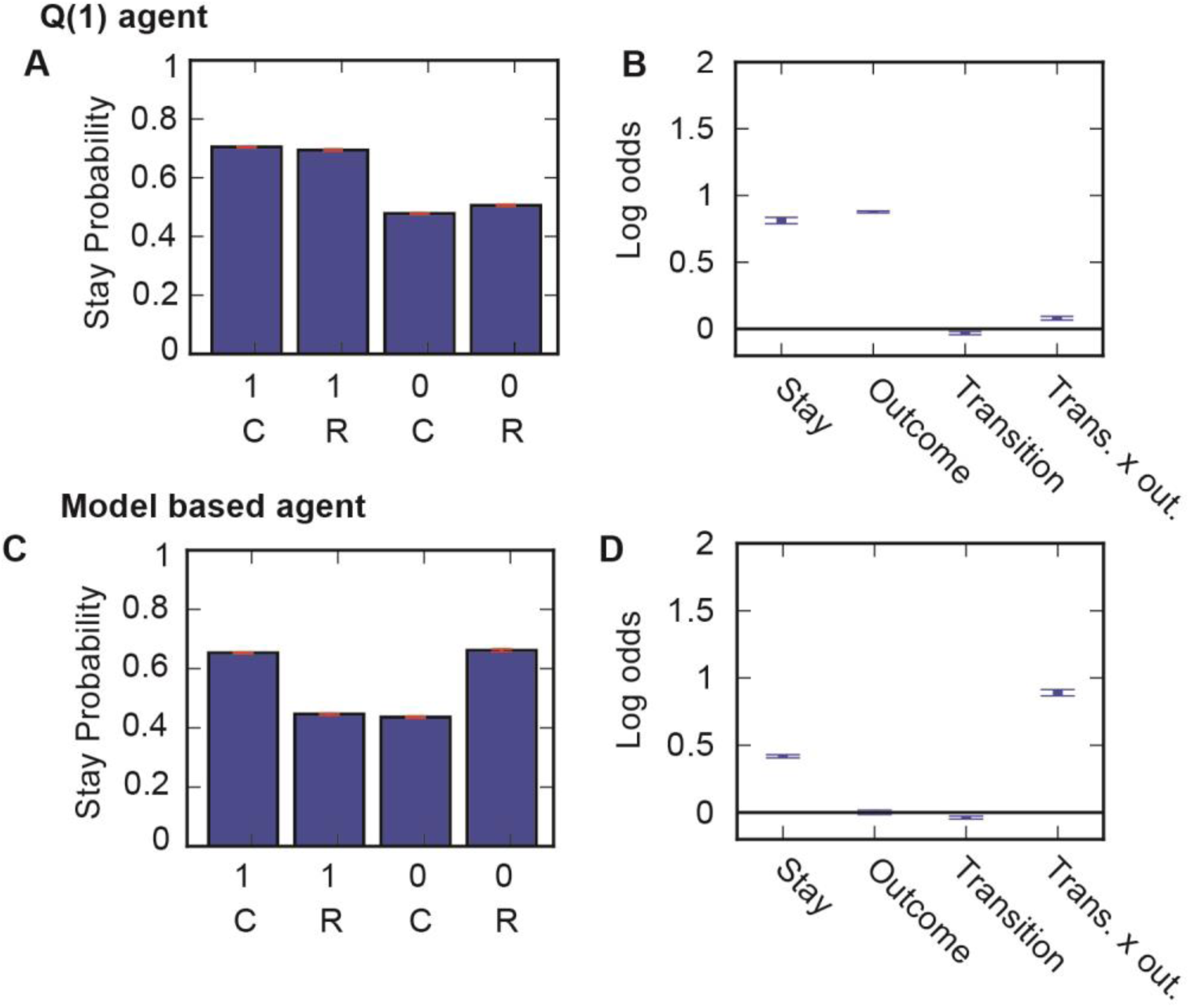
Original version of task. Behaviour of *Q*(1) (**A**, **B**) and model-based (**C**, **D**) agent with the same parameter values used in figure 3, simulated on the version of the two-step task used in Daw et al. 2011. (**A**,**C**) Stay probability analysis, (**B**,**D**) Logistic regression analysis using stay, outcome, transition and transition-outcome interaction predictors.

